# Precision of sensory representations in cortex is indexed by alpha power and pupil dilation, and tracks the dynamics of correlated noise

**DOI:** 10.64898/2026.11.18.767508

**Authors:** Sammi R. Chekroud, Nils Kolling, Jill X. O’Reilly

## Abstract

Sensory representations fluctuate in precision from moment to moment, shaping the fidelity of perception and memory. Population-coding theory proposes that these fluctuations are constrained by correlated variability within neural populations, yet linking these mechanisms to behaviour and brain state in humans has remained challenging. Here we combined EEG with an encoding–decoding framework to reconstruct orientation-selective population responses and quantify representational precision dynamically over time. Precision predicted subsequent working-memory performance and evolved in close correspondence with the dynamics of correlated variability between decoded orientation channels. Following stimulus onset, correlated variability was selectively suppressed among similarly tuned channels, the pattern predicted to maximise coding fidelity. Prestimulus alpha power and pupil size independently predicted representational precision, linking these dynamics to endogenous cortical and neuromodulatory state. Together, these findings identify candidate mechanisms and accessible physiological markers of representational precision in humans.

---

Engagement with sensory input fluctuates over time, even during simple tasks. During periods of high engagement, incoming stimuli are processed with high fidelity, whereas during disengagement, behaviour becomes more reliant on prior expectations and habitual responses (Ashwood et al., 2022; Honig et al., 2020). Endogenous fluctuations in arousal and attention occur over timescales of seconds to minutes and are closely linked to variability in behavioural performance (Gaillard et al., 2021). Mechanistically, what determines the precision with which sensory information is represented in the brain, and why does it vary over time?

At the level of neural populations, the precision of the information in a population of neurons is fundamentally limited by the correlation between firing units (Hazon et al., 2022; Zohary et al., 1994). In particular, noise correlations—shared trial-to-trial variability between units—place a key limit on how accurately information can be encoded (Cohen & Maunsell, 2009). Unlike independent noise, correlated variability cannot be averaged out across a population (Britten et al., 1992; Zohary et al., 1994; Ruff & Cohen, 2014). Noise correlations are reduced when observers direct attention towards specific stimuli or stimulus dimensions (Cohen & Maunsell, 2009, 2010; Ni et al., 2018), driving improved precision of sensory processing. Correlated variability between similarly-tuned neurons is proposed to be particularly detrimental to the precision of neural representations (Averbeck et al., 2006). Theoretical and empirical work has therefore highlighted reductions in noise correlations, particularly between units with similar tuning properties, as a principal mechanism by which the precision of population codes is controlled.

While such mechanisms are well established in invasive recordings, linking them to non-invasive measures of human brain activity remains challenging. We cannot directly observe correlations between individual neurons using EEG or fMRI, raising the question of how fluctuations in population-level precision might be tracked in humans. Whilst modelling approaches have recently been developed to estimate representational precision from neuroimaging data, these typically require large datasets and complex modelling approaches (Myers et al., 2015; Van Bergen et al., 2015), limiting their applicability in many research settings, including clinical studies. This limitation is particularly important given growing evidence that aberrant precision-weighting and population coding may underlie neuropsychiatric conditions such as psychosis (Fletcher & Frith, 2009; Sterzer et al., 2018).

Two candidate biomarkers of attentional and arousal state may provide indirect access to these mechanisms. Oscillatory activity in the alpha band is suppressed during states of high attention and reflects fluctuations in cortical excitability and coordinated population activity (Foxe et al., 1998; Haegens et al., 2011). Pupil size, in turn, tracks activity in neuromodulatory systems that regulate cortical state and signal-to-noise ratios (Aston-Jones & Cohen, 2005; Joshi et al., 2016; Reimer et al., 2016). Together, these measures offer a potential window into the neural conditions that shape population coding.

Here, we address three key questions. First, can the precision of sensory representations be estimated on a trial-by-trial basis from non-invasive EEG recordings, such that variability in measured neural precision can predict behavioural performance? Second, can fluctuations in precision be explained by changes in the structure of noise correlations? Third, can these changes be indexed by pre-stimulus biomarkers such as alpha power and pupil size?

We test the hypothesis that spontaneous fluctuations in task engagement modulate the precision of sensory representations by altering the structure of noise correlations in cortex, and further, relate these fluctuations in precision to readily measured biomarkers. To do this, we combine EEG recordings with an encoding–decoding approach that reconstructs orientation-selective population responses over time. This allows us to estimate, on a trial-by-trial basis, the precision of sensory representations and relate it to both behavioural performance and pre-stimulus biomarkers of attentional state. As predicted, during periods of high neural precision, noise correlations between similarly-tuned populations were decreased, precisely the change in noise structure that theory predicts would improve precision. Critically, we extend this framework to characterise the temporal dynamics of correlations between decoded feature-selective channels, providing a proxy for the underlying structure of population activity.

## Results

The dataset used here comes from a previously published study that addressed different research questions, beyond the scope of this paper (Chekroud et al., 2025). The task is illustrated in Figure 1a.

**Figure 1.**
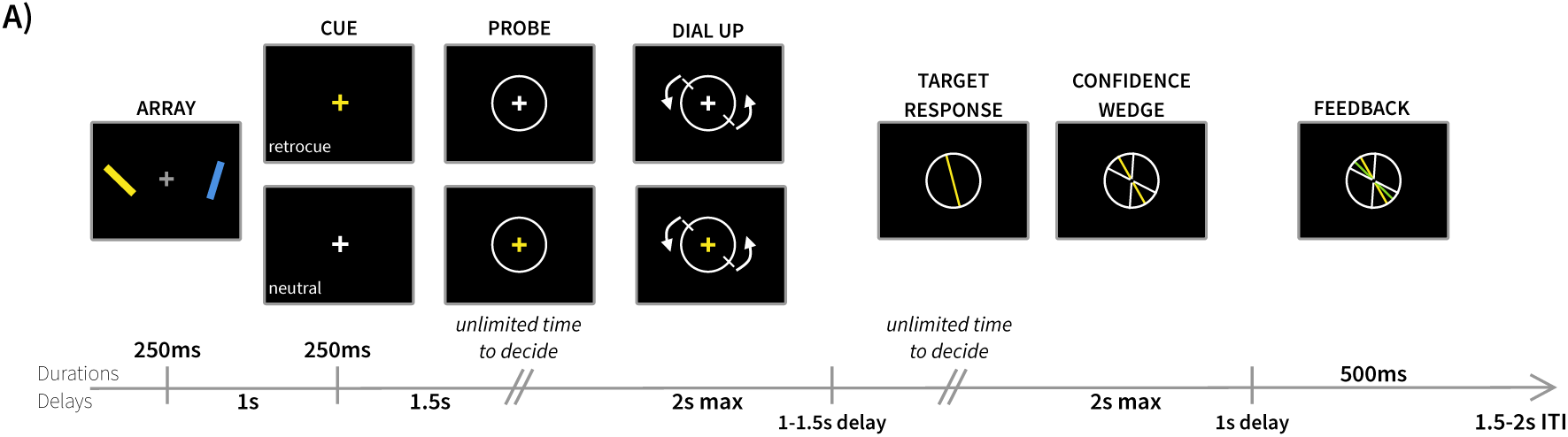
- Task schematic reproduced with consent from Chekroud et al., (2025) showing overall structure of a two-item working memory task where participants were required to reproduce the exact orientation of a previously encoded oriented bar.

In summary, participants performed a task in which two oriented line segments were held in working memory. On each trial, two line segments were presented, one in the left visual field and the other in the right. About 2.5s later, orientation was reported on a continuous scale (by rotating a sample line), giving a continuous measure of the precision of the representation, at the end of the working memory interval. Only one of the two line segments was probed for recall, allowing the relationship between recall precision and neural representation for the relevant (probed) and irrelevant (un-probed/distractor) stimulus to be probed separately. For reasons beyond the scope of the current analysis, on 50% of trials participants received a retrospective cue (“retro cue”) part way through the working memory interval that indicated which stimulus would be probed; this occurred one second after the stimuli were presented, so well after the time window analysed here.

### Time-resolved decoding reveals dynamic changes in sensory precision

To quantify the fidelity of sensory representations over time, we used an encoding-decoding model to reconstruct orientation-selective population responses using posterior EEG sensor activity (Fig 2A-C).

**Figure 2.**
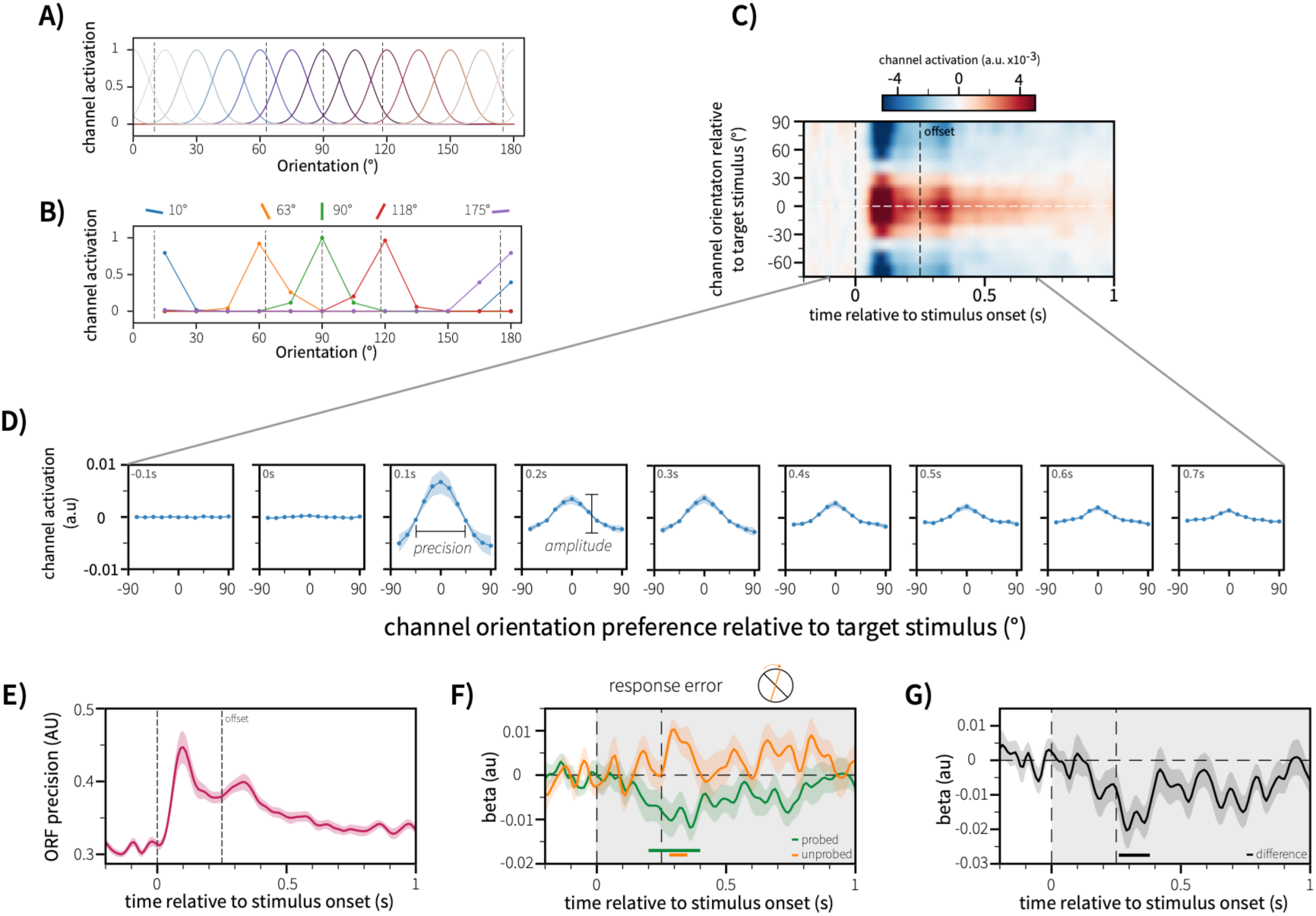
-Outline of orientation decoding and tuning curve decomposition approach. A) Activation profile of half-wave rectified cosine basis sets used to simulate tuned orientation channels. B) Simulated activation of orientation channels given stimulus orientations. Orientation channel activations are used to model the observed EEG sensor data. C) Ridge regression orientation decoding estimates decoded channel activations for each trial, reconstructing the underlying population tuning curve across time. Orientation response function is centred on the presented stimulus to visualise a stimulus-centred tuning curve across time. D) Orientation channel activations are collapsed over time bins of 50ms either side of the time point shown in the top-lek of each sub-panel. Schematics highlighting the tuning curve parameters estimated are shown. Precision and amplitude reflect the quality and amount of orientation-specific information available in the orientation channel activations. E) Stimulus onset is associated with rapid increases in the precision of sensory information. F) Response error is predicted by the precision of sensory encoding across trials. Precision of the item reported on each trial is negatively associated with recall error, reflecting that high precision of the probed item is associated with reduced error later in the trial (green line). Conversely, higher precision of the <u>unprobed</u> item is predictive of worse recall error (orange) later in the trial. G) The relationship between item-specific precision and working-memory error is significantly diWerent for the probed and unprobed item (black line shows the average within-participant diWerence), highlighting the specific nature of the relationship between item-specific precision and later memory performance.

This approach estimates the activation of latent orientation-tuned neuronal populations that would give rise to the observed EEG signal. The model learns a weight matrix describing the relationship between EEG sensor activity and orientation channel activation, then uses this mapping to decode the orientation-channel activity that best accounts for the held-out (test) EEG sensor data. The model was fit using 10-fold cross-validated ridge regression, yielding a trial-wise estimate of the orientation response function at each timepoint (see *Orientation Decoding* in *Methods*).

We then decomposed each orientation response function into two parameters: precision, reflecting the sharpness of orientation tuning, and amplitude, reflecting the strength of orientation-selective modulation (Fig 2D; see *Tuning Curve Decomposition* in *Methods*).

Stimulus onset was followed by a rapid increase in representational precision (Fig 2E). We observed two distinct peaks in the orientation response function – an early peak 100-200ms after stimulus onset, coincident with the P1/N1 complex, and a later peak at 300-400ms after stimulus onset, overlapping the P300. We hypothesised that the early peak reflects sensory encoding whilst the later peak may reflect transfer to working memory. A similar temporal profile was observed for the amplitude of the orientation response function (see *supplementary materials* for visualisation), indicating that stimulus onset increased both the quality (precision) and strength (amplitude) of orientation-selective information across visual cortical sensors.

### Neural precision at encoding predicts later recall fidelity

Previous research suggests that the precision of stimulus encoding varies from trial to trial (Fougnie et al., 2012), and gates the quality of subsequent recall from memory. We therefore asked whether trialwise fluctuations in the precision of the orientation response function predicted subsequent behavioural performance.

Using a time-resolved general linear model, we found that neural precision of the subsequently-probed item predicted recall error several seconds later, as measured by the orientation reproduction task. Specifically, higher precision in the period 200-400ms after stimulus onset was significantly associated with lower working-memory error (Fig 2F, green time-series), indicating that trial-wise variability in early sensory precision during encoding predicts the fidelity of subsequent memory-guided behaviour.

### Neural precision predicts recall fidelity in an item-specific manner

We next used the *unprobed* item on each trial to test whether the relationship between neural precision and performance was item-specific, or simply reflected some global state of improved encoding, such as arousal or attention. If the relationship between precision and behaviour simply reflected a global low-noise state, then higher representational precision of *all* visual stimuli would predict lower working-memory error. Alternatively, if behavioural performance is dependent on item-specific precision, then precision of the unprobed item should either be unrelated to behavioural performance, or interfere with it.

We found that increased neural precision for the unprobed item was associated with worse performance (increased response error) on the probed item (Fig 2F, orange timeseries; significant cluster 280-350ms post stimulus onset). The effects of precision of the probed and unprobed items differed significantly (Fig 2G, significant cluster 260-380ms post stimulus onset), indicating that the relationship between neural precision and recall fidelity is item specific and does not simply represent a generic high or low noise state.

Notably, the detrimental effect of the unprobed item emerged after stimulus offset, during the second peak in the orientation response function, suggesting some post-encoding interference between items when already encoded with high precision.

### Noise correlation suppression as a mechanism for increased representational precision

Having established that our measure of neural precision predicts behavioural performance, we next asked what neural mechanism could underlie these dynamic changes in representational fidelity. Population-coding theory suggests that correlated variability between neuronal populations limits fidelity of population codes, particularly when shared variability occurs between similarly-tuned populations (Averbeck et al., 2006). We therefore asked whether decoded orientation channels exhibited structured patterns of correlated variability consistent with empirical and theoretical work, and whether stimulus onset affected noise correlations as a function of tuning dissimilarity.

We calculated pairwise noise correlations between orientation-selective channels across time as a function of tuning dissimilarity (Fig 3A-C). Because ridge regularisation alters the covariance structure between channels, noise correlations were estimated using an unbiased OLS implementation of the decoding model (see *Noise Correlations* in *Methods* for more detail). Noise correlations were baselined relative to the pre-stimulus period.

**Figure 3.**
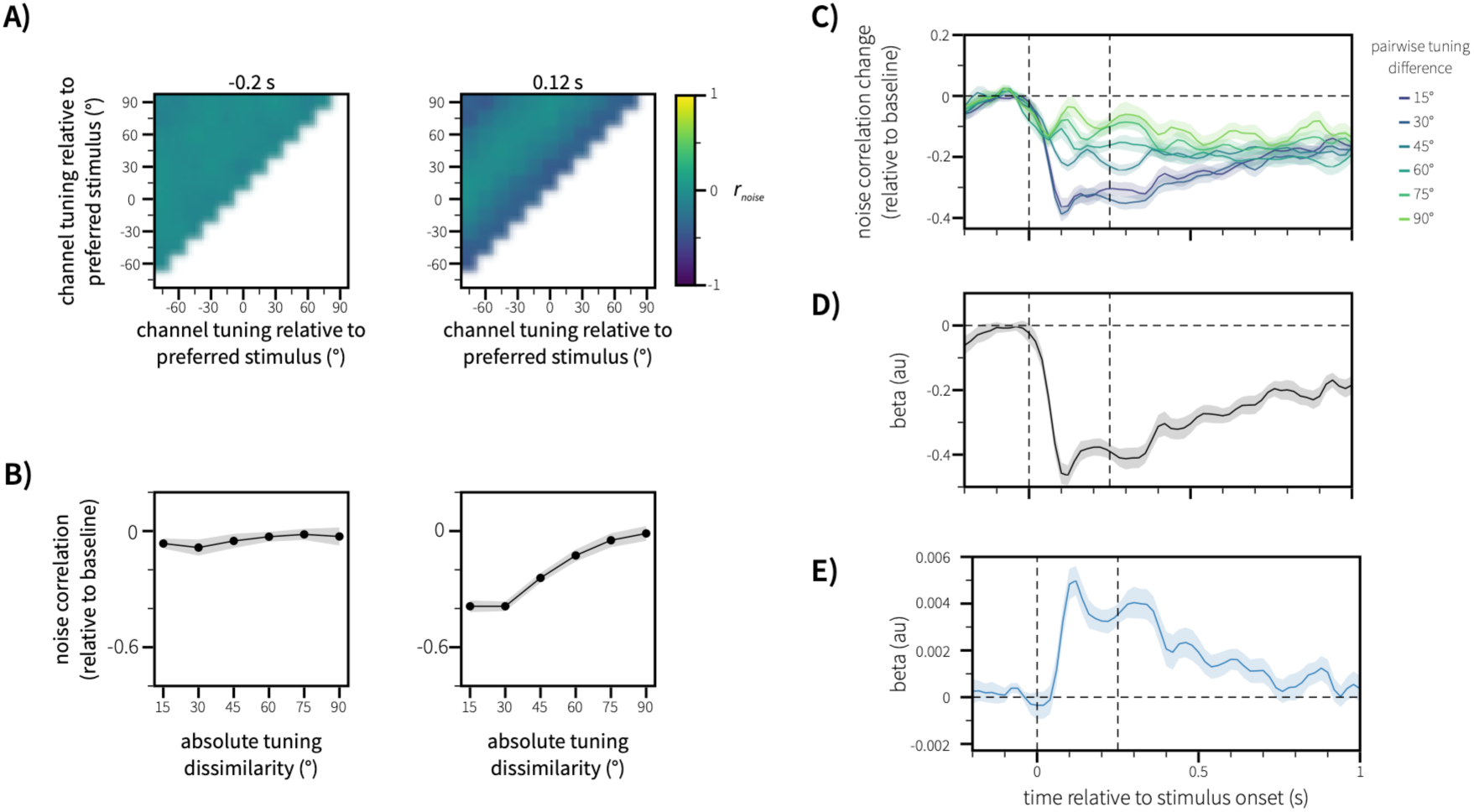
- Noise correlations between orientation channels. A) Pairwise noise correlations are calculated in steps of 20ms (±10ms), giving rise to a matrix of noise correlations between orientation channels across time, separately for each item in the stimulus array. The noise correlation matrix is baselined relative to the average noise correlation in the period 100ms to 0ms prior to stimulus onset. The noise correlation matrix is shown for two distinct time periods: during the pre-stimulus period and during active sensory processing (-0.2s and 0.12s relative to stimulus onset, respectively). B) Baselined noise correlations are visualised as a function of the unsigned pairwise tuning diWerence between orientation channels. C) Pairwise noise correlations rapidly decrease aker stimulus onset for all levels of tuning dissimilarity. Pairwise noise correlations are modelled as a linear combination of the mean noise correlation and pairwise tuning dissimilarity (in degrees). This revealed both a strong mean suppression of noise correlations for all orientation channels (D), and a positive association between orientation and noise correlation suppression (E). This highlighted that pairwise noise correlations were relatively more suppressed for similarly-tuned channels (dark lines in C) compared to orthogonally-tuned channels (light green lines in C)

Stimulus onset was followed by a rapid suppression of mean noise correlations across orientation-tuned channels (Fig 3D). Critically, suppression was strongest among similarly-tuned channels (Fig 3E, also visualised in Fig 3C) – the pattern predicted to improve the precision of the population code (Averbeck et al., 2006).

### Noise-correlation suppression predicts representational precision across individuals

Optimal coding theories predict that reductions in correlated variability should improve the precision of population codes. However, because the noise correlations are defined across repeated stimulus presentations, they cannot be estimated at the single-trial level. We therefore turned to cross-participant analysis to test the relationship between these measures. We predicted that participants showing stronger noise correlation suppression, particularly between similarly-tuned channels, should show greater precision in the orientation response function.

We first tested the across-participant relationship between mean noise correlation suppression across all channels (figure 4A, black line) and the mean precision of the orientation response function (Fig 4a, dark pink line) at each time point. Across participants, stronger mean noise correlation suppression was related to higher sensory precision (Figure 4C). Dynamically, these correlations were observed around the peaks of both precision and noise correlation suppression (around 100-200ms and 300-40ms after stimulus onset).

**Figure 4.**
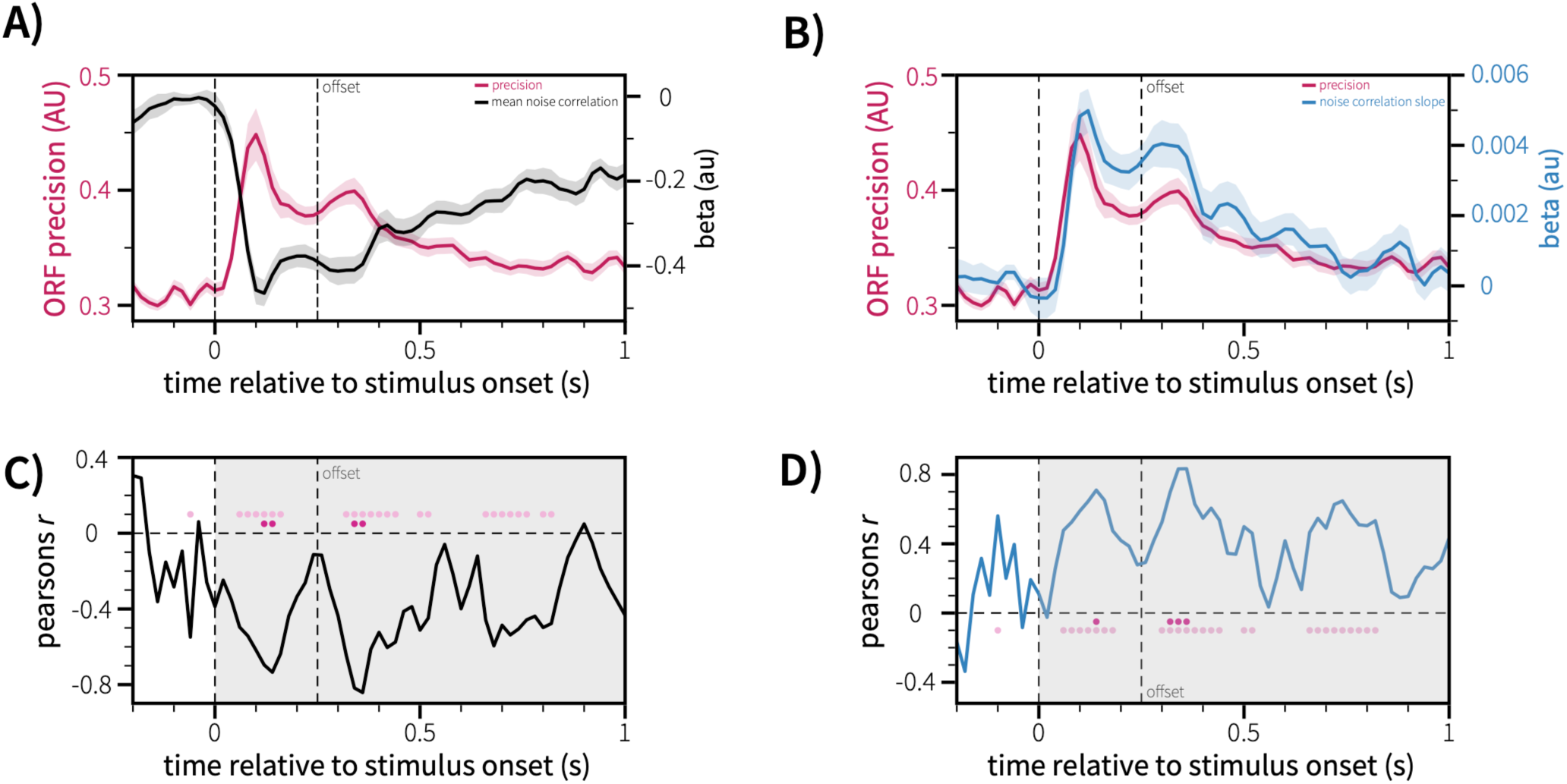
- A) reproduced from Figures 2 and 3, mean precision (dark pink) and mean noise correlation suppression (black) are overlaid for visualisation. B) reproduced from Figures 2 and 3, mean precision (dark pink) and the average slope of noise correlation suppression across orientation (blue) are overlaid for visualisation. Solid lines reflect the mean across participants, shaded regions represent the standard error of the mean. C) Across-participants, per-timepoint mean suppression of pairwise noise correlations is associated with higher precision during sensory processing. Negative correlations reflect that stronger noise correlation suppression was associated with higher precision across participants. D) Across-participants, per-timepoint, the slope of pairwise noise correlation suppression is associated with higher precision during sensory processing. Positive correlations reflect that a stronger slope for orientation (stronger suppression of similarly-tuned units vs orthogonally tuned units) was associated with higher representational precision across participants. Significant timepoints are marked with coloured dots. Timepoints that reached uncorrected-significance are marked with a light pink dot. Timepoints that survive Bonferroni correction (for the number of timepoints in the post-stimulus window marked by the shaded span along the x-axis) are marked in red.

We then tested the specific prediction that suppression of noise correlations between similarly-tuned neuronal populations should be particularly associated with increased precision. Across participants, stronger tuning-dependent suppression of noise correlations – indexed by the slope parameter of our noise correlation model (Fig 4B, blue line) – was associated with higher precision of the orientation response function (Fig 4B, dark pink line). Again, this relationship was observed around the peaks of both precision and noise correlation suppression (Fig 4D).

Together, these findings suggest that stronger suppression of noise correlations was associated with greater representational precision, particularly when noise correlations were preferentially suppressed between similarly-tuned channels, consistent with noise correlations as a candidate mechanism for controlling the precision of sensory representations.

### Accessible biomarkers of sensory precision

Using the analysis pipeline described so far, we are able to probe the precision of sensory representations using non-invasive EEG in humans, and their underlying mechanism (noise correlations). The precision of sensory representations is of interest in applied research – for example, ‘aberrant salience’ or excessive precision of sensory representation is proposed as a key mechanism is psychosis (Fletcher & Frith, 2009). In this context, paradigms requiring hours of focussed sensory tasks and complex pipelines are unsuitable.

We hypothesised that two readily measured indices of cortical state, namely alpha power and pupil dilation, could act as biomarkers for these computationally specific changes in cortical function (see *discussion* for background). We therefore asked whether precision of the orientation response function, on a trial-to-trial basis, is associated with systematic changes in alpha power and pupil size.

### Lower prestimulus alpha power predicts increased representational precision

Using a time-resolved general linear model, we found that lower baseline alpha (which is associated with higher arousal), measured in the period 500ms before stimulus presentation, was associated with higher precision during sensory processing (significant cluster 60-190ms post stimulus presentation, figure 5A).

**Figure 5.**
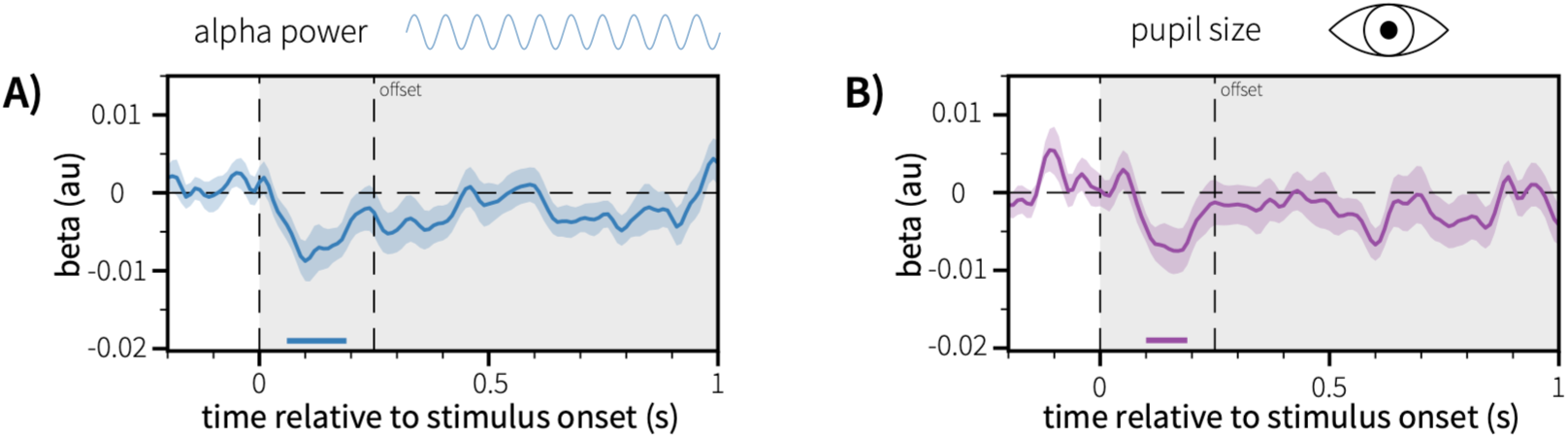
- Baseline markers of attentional state are predictive of the average precision of sensory information across items in the stimulus array. Baseline alpha power (A) and baseline pupil size (B) in the 500ms prior to stimulus are negatively predictive of representational. Low baseline alpha power and a more constricted baseline pupil size are associated with increased representational precision during early sensory encoding. Grey shaded areas on axes highlight the time-window used for cluster permutation testing. Horizontal lines above and parallel to the x-axis denote time periods of significance following cluster permutation testing. Solid lines reflect the across-participant mean, shaded ribbons reflect the standard error of the mean.

### Smaller pre-stimulus pupil size predicts increased representational precision

Pre-stimulus pupil size was also associated with representational precision: Smaller pupil size predicted greater representational precision during sensory processing (statistically significant 100-190ms post stimulus onset, figure 5B). In addition, smaller pupil size was associated with and decreased pre-stimulus alpha power at the single-trial level (*t*(8260.456) = 3.50, p = 0.000467).

Intriguingly, although we observed a triangle of bilateral relationships (pupil and alpha predict neural precision, and pupil predicts alpha), a formal mediation analysis showed no evidence that the effect of either biomarker on neural precision is mediated by the other (see *supplementary information*). This suggests that alpha power and pupil size may provide partially independent indices of the neural states associated with changes in representational precision.

Notably, the direction of the pupil effect differs from studies linking larger pupil size to increased cognitive effort or arousal. In the present task, however, behavioural performance depended on the precision of sensory representations rather than response vigour or sustained cognitive effort. Under this interpretation, smaller pupil size may reflect cortical states associated with reduced variability and more precise sensory encoding, consistent with previous work linking pupil constriction to reduced variability in perceptual decision-making (Murphy et al., 2014).

## Discussion

Endogenous fluctuations in brain state shape the fidelity of sensory representations in cortex. Using time-resolved EEG and an encoding–decoding model of orientation-tuned channels, we show that spontaneous variation in pre-stimulus alpha power and pupil size predicts the precision with which visual information is encoded, which in turn predicts subsequent behavioural performance. Critically, increases in representational precision are accompanied by decreases in noise correlations between feature-selective populations, particularly among similarly tuned channels. These findings link behavioural variability, large-scale brain signals, and population coding, suggesting that fluctuations in task engagement modulate sensory processing by altering the correlational structure of population activity.

Our findings can be naturally interpreted within theoretical frameworks that emphasise the role of precision in perceptual inference, such as predictive coding accounts(Friston & Kiebel, 2009; Rao & Ballard, 1999). In these models, the influence of sensory input on downstream processing depends on its precision relative to prior expectations. However, the neural mechanisms that determine this precision remain underspecified. The present results suggest that changes in the correlational structure of population activity may provide one such mechanism. Reductions in noise correlations—particularly among similarly tuned units—would increase the fidelity of sensory representations (Averbeck et al., 2006), effectively increasing the precision of bottom-up signals. Conversely, elevated correlated variability may reduce the impact of sensory input, biasing processing towards internally generated expectations. From this perspective, fluctuations in alpha power and pupil-linked arousal may index shifts in the balance between externally driven and internally driven processing by modulating the structure of population codes. This interpretation is consistent with proposals that aberrant precision-weighting contributes to perceptual and belief disturbances in neuropsychiatric conditions such as psychosis (Fletcher & Frith, 2009; Sterzer et al., 2018), and raises the possibility that alterations in population-level correlations may underlie these effects.

Consistent with optimal coding theories, we found that noise correlations between decoded orientation channels were substantially reduced following stimulus onset, in parallel with increases in sensory precision. Moreover, noise correlations were most strongly attenuated among similarly tuned channels, where correlated variability is most detrimental to population coding (Averbeck et al., 2006). The noise correlation account of sensory precision is in contrast to classical gain theories, which propose that attention selectively enhances the responsiveness of neurons tuned to attended features (Treue & Martínez Trujillo, 1999). Such models account well for feature-specific enhancement, but not for endogenous fluctuations in precision that occur prior to stimulus processing. In contrast, the control of noise correlations provides a global mechanism for controlling the fidelity of sensory representations (Ni et al., 2018), and accounts for a large portion of attentional enhancement in sensory processing (Cohen & Maunsell, 2009).

While related, our measures of precision and noise correlation capture distinct aspects of population coding. Precision reflects the sharpness of the decoded tuning function at the single-trial level, whereas noise correlations quantify shared variability across trials between channels. Although reductions in noise correlations are expected to increase precision, and noise correlation and precision metrics correlate across individuals, they are not equivalent measures, and multiple mechanisms may contribute to changes in precision. Our findings therefore support a link between these quantities, rather than a one-to-one correspondence.

A noteworthy aspect of our approach is the ability to estimate the precision of decoded population responses independently of the amplitude, allowing us to show that behavioural performance is specifically linked to precision rather than overall signal strength. In contrast, previous research has largely investigated the relevance of the overall strength (amplitude) of the decoded information content (Myers et al., 2015; van Ede et al., 2018; Wolff et al., 2017).

We found that fluctuations in alpha power and pupil size track changes in the precision of sensory representations. Beyond their practical value as accessible biomarkers, these relationships point to a potential mechanism by which global brain state influences the correlational structure of population activity.

Alpha power and pupil size covaried at the single-trial level and showed convergent relationships with sensory precision over time. While these measures are often interpreted as reflecting distinct processes—alpha power indexing cortical excitability and local coordination, and pupil size indexing global neuromodulatory tone—their shared relationship with representational precision suggests a common underlying influence on population-level activity. One possibility is that both measures reflect changes in the balance between coordinated and independent neural firing, thereby modulating the structure of noise correlations.

Alpha oscillations in sensory cortex are thought to reflect rhythmic modulation of neuronal excitability at the population level (Haegens et al., 2011). Higher alpha power has been associated with reduced firing rates and increased phase-locking, consistent with more coordinated—and therefore more correlated—neural activity. Conversely, reductions in alpha power are associated with more asynchronous firing, which may reduce correlated variability and enhance the precision of population codes. Consistent with this account, anticipatory reductions in alpha power induced by spatial cues have been shown to enhance stimulus decoding activity (Griffiths et al., 2019; van Ede et al., 2018). Our results extend these findings by showing that *endogenous* fluctuations in alpha power similarly predict the precision of sensory representations, even in the absence of explicit attentional cues.

Pupil size provides a complementary index of global brain state, tracking activity in neuromodulatory systems. Noradrenergic activity from the locus coeruleus has been closely linked to pupil dilation (Aston-Jones & Cohen, 2005; Gilzenrat et al., 2010; Joshi et al., 2016; Disney, 2021; Lloyd et al., 2023), while cholinergic modulation has been implicated in attentional control and sensory processing (Disney et al., 2007). In particular, manipulating cholinergic activity has been shown to causally change the signal-to-noise ratio of sensory cortical neurons and reduce noise correlations (Herrero et al., 2013; Minces et al., 2017), providing a plausible mechanism linking neuromodulation to the effects observed here. While pupil size is more commonly associated with noradrenergic activity, its relationship with cholinergic tone is less clear, and both systems are likely to contribute to the observed effects.

Taken together, these findings suggest that alpha power and pupil size provide complementary, but not identical, indices of underlying neural states that shape the precision of population codes. Importantly, our results suggest a link between these macroscopic biomarkers and the structure of population activity that determines sensory precision. While we cannot directly measure noise correlations at the single-trial level, both alpha power and pupil-linked arousal have been independently associated with changes in coordinated neuronal activity and cortical state. In the present study, lower pre-stimulus alpha power and smaller pupil size predicted higher sensory precision, while, at the population level, increased precision was associated with reduced noise correlations—particularly among similarly tuned channels. Taken together, this pattern is consistent with the interpretation that fluctuations in these biomarkers reflect changes in the degree of shared variability within neural populations. Under this account, states characterised by reduced alpha power and altered neuromodulatory tone may promote more independent neuronal activity, thereby reducing correlated noise and enhancing the fidelity of sensory representations.

Several limitations should be considered when interpreting these findings. First, our estimates of population activity and noise correlations are derived from decoded feature-selective channels rather than directly recorded neurons. While this approach captures population-level structure, it necessarily depends on assumptions about the mapping between sensor-level signals and underlying neural activity, and may be influenced by spatial mixing and model specification. Second, although we link fluctuations in alpha power and pupil size to changes in representational precision, these measures, especially pupil size, provide indirect indices of brain state and are subject to multiple driving influences, therefore a correlative approach alone does not allow us to isolate the specific neuromodulatory or circuit-level mechanisms involved.

Despite these constraints, the convergence of behavioural, physiological, and population-level measures suggests that endogenous fluctuations in brain state shape sensory processing by modulating correlated variability, thereby determining the precision of neural representations.

## Methods

The current article presents a re-analysis of data from a previous study, with a novel analytical pipeline to investigate distinct research questions. The original data were previously published, focused on how visual selective attention during working-memory maintenance (using retrocues) affects metacognitive judgements (Chekroud et al., 2025). Data are freely available on OSF (DOI: 10.17605/OSF.IO/98A3K). Here, we explore sensory encoding – the transformation of sensory information into an internal, mnemonic representation. We present a new analytical pipeline for the analysis of sensory encoding, parameterising the precision and amplitude of internally held representations.

### Participants

We analyse data from the final sample of 20 participants described in the original study (age range 18-36; 14 female; 1 left-handed). All participants reported normal or corrected-to-normal vision and not being colourblind. All participants provided written informed consent prior to participation, and were reimbursed £15/hour for their time. Experimental procedures were reviewed and approved by the Central University Research Ethics Committee of the University of Oxford.

### Task design

Participants performed a two-item visual working-memory task with a delayed orientation-reproduction response from memory. Trials began with an inter-trial interval drawn randomly between 1 and 2-seconds, with a central grey fixation cross presented throughout. Participants then encoded two oriented coloured bars (250 ms duration) presented either side of the fixation cross (centred 6° visual angle from the fixation cross). After a 1-second delay, the fixation cross changed colour for 250 ms to either match the colour of one of the encoded bars (retrocue, 50% of trials), or to white (neutral cue, 50% of trials). There was then a further 1.5-second delay before the probe stimulus appeared. The probe stimulus consisted of a central fixation cross surrounded by a circle (diameter 5.7° visual angle). On cued trials, where the fixation cross during the delay had changed colour to match one of the encoded items, the fixation cross at the probe was coloured white (making it imperative to process the cue). On neutral trials, the fixation cross during the probe display changed colour to match the to-be-reported item. Participants had unlimited time to decide on the orientation they wished to report, before pressing the ‘space’ bar (with their left hand) to indicate they were ready. Upon pressing the space bar, the response dial appeared – two white lines on opposite sides of the probe circle. The starting orientation of this response dial was generated randomly (drawn between 0 and 179°), and participants used the mouse to rotate the dial around the circle to replicate the probed orientation. Participants made a left mouse click to confirm their response. After their initial orientation reproduction, participants were required to make a response to indicate how confident they were in their response, before being given feedback that conveyed information about their response error and the accuracy of their metacognitive judgement. Further details of how this was generated can be found in the original publication (Experiment 2, Chekroud et al., 2025), as this is out of the scope of the current article and unrelated to our question of interest.

### Pupillometry acquisition and preprocessing

Pupillometry data were collected using an EyeLink 1000 (SR Research) with the participants’ head resting on a chin rest. During the task, gaze and pupil size were continuously tracked for both eyes simultaneously, at a sampling rate of 1000 Hz. Prior to acquisition, the eyetracker was calibrated using built-in calibration and validation routines from the EyeLink software.

Eyetracking data were parsed and preprocessed separately for each block of task, per participant, using custom code in python version 3.13.5. Data were first converted from the edf to asc format, before being parsed. Where possible, we used pupil data from both eyes. Where blocks of task data were especially noisy for individual eyes (for example, the eyetracker lost sight of one eye, or a change in head position moved one eye out of focus), we considered only the eye with the least noise. Where recording of both eyes was lost for a block, the entire block was removed from subsequent analysis.

Blinks were identified using an automated routine based on the first derivative of the pupil signal and missing data due to blink periods. We corrected blink periods with a ±75 ms window around the blink to account for corrupted data, using a linear interpolation over the blink window. The pupil timecourse for each block was smoothed using a gaussian smoothing kernel with a 50ms standard deviation blur. To account for slow drifts in pupil size in each block, unrelated to the task (for example, pupil size is larger at the beginning of a task block compared to the end), we estimated a cubic polynomial fit to the pupil timecourse and subtracted this from the signal. Where pupil data in both eyes was suitable, we averaged pupil size across both eyes to give a single time series for analysis. Finally, data were epoched between-0.5 and 1.25 seconds relative to the stimulus array onset for further analysis. For statistical analysis, we only considered trials where less than 30% of the stimulus epoch was contaminated by blinks

### EEG acquisition and preprocessing

EEG data were collected using a SynAmpsRT amplifier and Curry 8 Software (Compumedics) using a 61-electrode array following the international 10-10 system. Data were reference to the left mastoid during recording, and re-referenced to an average-mastoid reference offline. The ground was placed on the upper left arm, and two bipolar electrooculograms (EOG) were recorded. One pair was placed above and below the left eye to measure the vertical EOG and identify blinks, the other placed lateral to each eye to record the horizontal EOG and identify saccades. Data were sampled at 1000 Hz during acquisition and stored for analysis.

Preprocessing was performed separately for each task session for each participant. EEG data were imported and preprocessed using the MNE-python toolbox version 1.90 (Gramfort et al., 2013) and custom code. Raw data were bandpass filtered between 1 and 40 Hz, and bad channels identified during recording were interpolated. Data were then epoched between-1 and +1.25 seconds relative to the onset of the array stimulus. Denoising was performed on the epoched data using the ‘infomax’ routine to all sensors. Components related to eye movements were detected by correlation with the horizontal and vertical EOG, and subsequently removed from the data. Trials were removed based on the within-trial variance using an automated procedure that looks at signal variance at a significance threshold of 0.05 (generalised studentized deviate test, as in e.g. Chekroud et al., 2025). Trials were subsequently visually inspected to identify any remaining trials containing excessive noise, to be removed from analysis.

### Time-frequency decomposition

Spectral power was estimated at frequencies using a multitaper decomposition. For the stimulus epoch, data were downsampled to 500 Hz and single-trial power was estimated for frequencies between 1 and 30 Hz. For each frequency, we used a fixed 300 ms time window such that the number of cycles changed with estimated frequency.

For analyses relating to single-trial alpha power, we considered only spectral power in the 17 most posterior channels (P7, P5, P3, P1, Pz, P2, P4, P6, P8, PO7, PO3, POz, PO4, PO8, O1, Oz, O2). Data were averaged across frequencies in the alpha (8-14 Hz) range, before averaging across these posterior sensors to get a single-trial timecourse of alpha power.

### Orientation decoding

We quantified the information contained in recorded scalp activity using a population tuning curve model to decode orientation information from the multivariate scalp pattern across posterior (primarily visual) EEG sensors. In this model, activity across sensors is modelled as a linearly-weighted combination of activation of a set of theoretical, orientation-tuned channels (or neuronal populations). A weight matrix is learned that describes the transformation between EEG sensor space and orientation-channel space, and applied to held-out test data to decode (or infer) the activation of orientation channels that best accounts for the observed EEG data. As a result, this approach infers the activation of the underlying cortical generators of orientation-specific activity in the observed EEG data.

This encoding-decoding approach (e.g. Myers et al., 2015; Van Bergen et al., 2015), unlike pattern classifiers or distance-based metric approaches (e.g. Wolff et al., 2017; van Ede et al., 2018), mirrors the underlying circuit-level computations in visual cortex that would give rise to the observed sensor data. This allows for a more interpretable inference about how those computations change under different neural conditions, such as varying levels of arousal. However, many of the results presented here can be reproduced using a (mathematically different) approached based on the Mahalanobis distances between patterns in representational/correlation space (see *supplementary information*).

The encoding-decoding model starts with the assumption that scalp activity relating to the orientation of a stimulus is related to the activity of a series of neuronal populations tuned to different orientations. We model stimulus orientation as the activation of a series of theoretical neuronal populations using weights from a linear basis set of population tuning curves. The activation of these orientation channels is then decoded from the evoked scalp activity, generating an orientation response function describing the orientation-specific information contained in the EEG across time.

Preprocessed epoched data were lightly smoothed using a gaussian kernel (20ms standard deviation) and the 17 most posterior visual sensors were selected for further analysis (electrodes P7, P5, P3, P1, Pz, P2, P4, P6, P8, PO7, PO3, POz, PO4, PO8, O1, Oz, O2). The scalp pattern across these sensors was demeaned at each time point, effectively removing global gain differences between timepoints that would be unrelated to the active processing of orientation. Separately, for each item in the encoding array, we implemented a machine-learning pipeline to decode the activation of our theoretical neuronal populations.

Firstly, we simulated the activation of our theoretical neuronal populations, *f(s)* using a half-wave rectified cosine function raised to the 15^th^ power

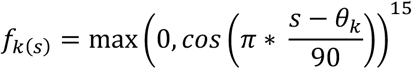

where θ_k_ is the preferred orientation of the *k*th *orientation channel*. We used 12 channels with orientation preferences equally spaced between 15-180° (as the stimuli in this experiment had 180 degree rotational symmetry). This resulted in two orientation channels maximally tuned to the vertical (90°) and horizontal (180°) axes. This creates an overlapping set of basis functions that describe the activation of theoretical, orientation-tuned neuronal populations given a particular stimulus orientation. Separately for each item presented in the array, we used these basis sets to simulate the activation of each orientation channel given the orientation of the stimulus presented on each trial.

Our decoding pipeline used a Ridge regression which predicted the orientation channel activities using the EEG sensor data. Previous research has used ordinary least squares (OLS) in these population tuning curve models where singular stimuli are presented on the screen (Myers et al., 2015). Firstly, as multiple oriented lines were presented in the stimulus array, activity across the scalp will reflect a combination of both stimuli. Ridge regularisation thus allows a data driven penalisation of uninformative scalp sensors when decoding orientation channel activations underlying individual stimulus orientations. For example, stimuli were presented in different hemispheres, and ridge penalisation can effectively down-weight the contribution of ipsilateral posterior sensors to the decoder if their trial-to-trial fluctuations did not covary with stimulus orientation.

Secondly, we aimed to increase our decoding sensitivity at the single-trial level. Ridge regularisation allows the model to take into account the *expected shape* of channel activations (based on their tuning) and optimise the model accordingly. The shrinkage of coefficients through application of an L2-norm penalty in ridge regression significantly decreases variance across channels (in this case, single-trial noise) while increasing model bias. In reducing the variance of channel activations, it effectively *smooths* the tuning curve as a function of orientation, which also aids in later estimation of tuning curve precision and amplitude. Overall, this ridge regularisation improves single-trial sensitivity. However, the selection of an appropriate amount of regularisation is essential by tuning the regularisation hyperparameter, *alpha,* to appropriately balance this bias-variance trade-off (see *hyperparameter tuning)*.

For estimating orientation channel activations on each trial, separately for each decoded item, we used a 10-fold cross validation. Sensor data were normalised across trials using the mean and variance of sensor activity in a pre-stimulus period between 500ms to -100ms relative to stimulus onset. Standardising sensor data at each time point can make the decoder sensitive to noise at individual time points, while standardising across the entire stimulus epoch would cause leakage of sensory information between time points and bias decoding. We chose this baseline period to normalise the sensor data as it should reflect variability across sensors unlinked to the to-be-decoded orientation, thus applying similar scaling across sensors for the ridge. The mean and variance were estimated on the within-fold training set, and the same values were applied to re-scale the test data. The ridge regression was fit, and estimated ridge weights were applied to derive decoded channel activations for the test data. This resulted in a trial-wise estimate of decoded channel activations describing the activation of orientation-tuned neuronal channels, reflecting the orientation information contained in the scalp EEG.

### Hyperparameter Tuning

We optimised the regularisation strength, *α,* independently for each item in the array, allowing item-specific penalisation of sensor contributions to the decoded orientation channel activations. To obtain a single, interpretable regularisation strength per participant (and item), we pooled sensor data across trials and timepoints within the stimulus window 0.1-0.6 s to form a training set of shape [N_samples*time_, N_sensors_] and the corresponding tiled design matrix of channel responses. This stimulus window was chosen to optimise *α* over a time-period where we expected there to be a strong orientation-specific signal in the evoked sensor data. We then performed grouped 5-fold cross-validation (grouped by trial) so that entire trials were held out in the test data, preventing leakage due to autocorrelation across timepoints within trials. Sensor data were standardised before model fit. *Alpha* was tuned by grid search over 50 logarithmically spaced values between 10^-3^ and 10^7^, encompassing a range from both effectively unregularized (analogous to OLS estimation) and strong coefficient shrinkage.

Model scoring in this grid search was performed per held-out sample using the cosine similarity between the decoded and model channel activations. Cosine similarity between two vectors is easily interpretable (1 when they point in the same direction in high-dimensional space, 0 when orthogonal, and-1 when they point in opposite directions) and it is scale invariant – thus it is sensitive to the direction of the two vectors, not the magnitude of their direction. Thus, it prioritises sensitivity to the shape of the predicted channel activations and is robust to changes in amplitude over time – critical, as we expected orientation-information amplitude to vary over time. Alternatively, one could use the mean-squared error (MSE) or per-sample correlation coefficient as scorers. MSE is a poor scorer in this instance as it fails to account for the expected shape of predicted channel activations and is sensitive to channel activation amplitudes. Optimising for per-sample Pearson’s correlation can result in over-regularisation and poor prediction. Pearson’s correlation subtracts out mean channel activations in its implicit demeaning. However, channel responses to orthogonally-tuned orientation channels are a feature of the data (channel activations are low when the stimulus orientation is far from its preferred orientation). Optimising for Pearson’s correlation can lead to flat channel activation predictions where there is a global offset in channel activations (i.e. where all channels are active and there is a cosine-shaped tuning profile on top of this offset).

We chose a global *α* (one *α* per participant and item, across all trials and timepoints) so that changes in channel activations reflect changes in underlying neural signal rather than arbitrary changes in regularisation. To avoid over-regularisation and selection of an α tuned to small, possibly noisy peaks in the cross-validation curve, we selected the smallest *α* whose mean cross-validated score reached at least 90% of the maximum improvement above baseline (where baseline is the model score at the smallest value of *α*). Where regularisation showed no improvement from the smallest assessed level of regularisation, we accepted numerically the best *α*. This 90%-of-max selection criteria also accounts for the asymmetric sampling of possible regularisation values, where excessive regularisation could result in increasingly small improvements in model score at the expense of introducing more bias. The selected item-specific global α was then used for trial- and timepoint-wise estimation of channel activations when estimating the orientation tuning curve.

### Tuning curve decomposition

The main goal of our analytical method is to better quantify the neural representation: how much information is contained in the signal, and what is its quality? Previous studies have taken a mahalanobis distance approach to stimulus decoding, convolving the mahalanobis distances with the cosine of their respective bin centre, then averaging across these convolved distances to create a single measure of stimulus information, or sensory evidence (van Ede et al., 2018; Wolff et al., 2017). This effectively estimates the amplitude of a fixed-width cosine function. Instead, we developed a two-stage process to decompose the distribution of channel activations and leverage the circularity of the stimulus feature space. In brief, we estimated the best-fitting cosine functions in two stages to best capture two cognitively and computationally meaningful parameters: the precision of the neural representation, and its amplitude.

For each item in the encoding array, and each time point, we took the vector of channel activations and minmax-scaled this, such that the activations occupied the range from 0 to 1 – the same value range for the cosine function. We then used a non-linear least-squares optimisation (using the default ‘Trust Region Reflective’ algorithm) to fit a flexible cosine model to the minmax-scaled orientation channel activations of the form:

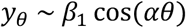

where *θ* is the angular distance (in radians) between each channel’s preferred orientation and the trial’s stimulus orientation, and *α* indexes the precision of the tuning curve (where larger values indicate narrower, more orientation-selective tuning). We also included a free-parameter for amplitude scaling *β*_1_ to accurately fit the precision parameter.

We used a non-linear optimisation rather than the more commonly used Nelder-Mead method with *fminsearch*, as this non-linear method allowed us to constrain parameter estimates. We constrained the precision parameter, *α*, between 0 and 3 to avoid extreme values that may be dominated by noise. Further, a negative value for the precision of the information is uninterpretable, as a precision of 0 would be represent a broad distribution of channel activations reflective of no orientation tuning in the signal.

The fitted precision parameter was then smoothed with a gaussian kernel (20ms standard deviation), and used to model tuning curve amplitude. Within each trial, *i,* the amplitude of the reconstructed channel activations at each timepoint, *t*, was estimated by modelling channel activations as a cosine function centred on the stimulus orientation. Specifically, channel activations *y_c_*_,*t*,*i*_, were modelled as:

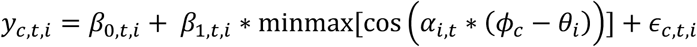

where *φ_c_* denotes each channel’s preferred orientation, *θ_c_* is the stimulus orientation on trial *i, and α_i_*_,*t*_ is the previously estimated precision parameter that controls the width of the cosine basis. The precision parameter at each trial and timepoint was constrained to have a minimum value of 0.001 – any lower and the design matrix is a vector of ones, and the model fails to converge. The cosine basis was minmax normalised to scale to the unit range before fitting. An intercept term, *β*_0_, was included to model baseline shifts in channel activations, such that *β*_1_ represents the relative modulation of orientation-selective activations above baseline, reflecting the *amplitude of the orientation-selective information* available in the tuning curve.

### Noise Correlations

We tested whether changes in underlying noise correlations between orientation-tuned channels could provide a mechanistic explanation for the changes in tuning curve precision we observed. In the cellular neuroscience literature, inter-neuronal correlations can take two primary forms: *signal correlation*, the correlation between two neurons to a range of stimuli (reflecting their tuning similarity), and *noise correlation*, reflecting the correlation between two neurons in response to repeated presentations of identical stimuli (Cohen & Kohn, 2011; Cohen & Maunsell, 2009). Correlated noise cannot be averaged out from the population response, and significantly degrades population codes (Britten et al., 1992; Zohary et al., 1994). Recent evidence has highlighted noise correlation reductions as a candidate mechanism by which attention operates on the precision of population codes (Cohen & Maunsell, 2009).

To calculate noise correlations in our data, we fit a distinct tuning curve model. Rather than using a ridge regression approach, we fit a standard OLS decoder pipeline to decode orientation channel activations. Ridge regression provides superior single-trial sensitivity in the tuning curve through regularisation. This regularisation reduces the across-channel variance in decoded channel activations, effectively smoothing the tuning curve and realising a more stable fit. This smoothing occurs as the ridge penalisation causes neighbouring channel activations to be more similar than orthogonal channel activations, artefactually increasing signal and noise correlations between orientation channels with similar tuning preferences. It also reduces the signal and noise correlations between orthogonally-tuned orientation channels. While this increases the signal available in the single-trial tuning curve (important when looking at single-trial relationships with physiological markers), it alters the across-channel covariance structure. So, to explore noise correlations, we re-implemented an OLS version of the pipeline as an *unbiased estimator* of channel activations to explore noise correlations, as channel activations are estimated independently of one another. Data were preprocessed as in the Ridge approach, but EEG sensors were not standardised prior to estimation of the weight matrix that describes the transformation from sensor space to orientation channel space.

Separately for each possible stimulus orientation, we calculated the Pearson’s correlation coefficient between orientation channels in the stimulus-centred tuning curves. This was only performed for stimulus orientations where there existed at least three trials with that orientation (the minimum needed to calculate a correlation). This gives rise to a 12 x 12 correlation matrix that describes the correlational structure between orientation channels for each possible stimulus orientation. Correlation coefficients were fisher-transformed then average across orientations to give one correlation matrix per item. This process was performed in steps of 20ms across the stimulus epoch, with a window of ±10 ms (we averaged the channel activations in this time window to reduce noise in the analysis), giving rise to a lightly-smoothed measure of noise correlations across time.

To analyse noise correlations, the derived correlation matrix was averaged across items in the stimulus array to give a metric of mean noise correlations over time. This noise correlation matrix was baselined relative to the average noise correlation in the period 100ms to 0ms prior to stimulus onset, to highlight stimulus-induced changes in noise correlations (raw noise correlations are presented in the *supplementary materials,* for completeness). We then unravelled this correlation matrix to give correlations as a function of tuning-similarity between orientation channels (15, 30, 45, 60, 75 and 90 degrees dissimilarity) to explore specific hypotheses about efficient population coding.

Optimal coding theories argue that limits on population codes are placed by correlations between similarly-tuned units with high signal correlation, where strong co-activation prevents the fine-grained disambiguation between incoming stimuli (Averbeck et al., 2006). When two orientation-tuned neurons have high signal correlation (similar orientation tuning) *and* high noise correlations (their endogenous variability across repetitions is correlated), a higher-order decoder is unable to reliably discern the stimulus feature based on their relative activation. Two orientation-tuned neurons with *low signal correlation* (orthogonally-tuned) are highly separable based on their activations due to the incoming sensory input (larger differences in firing rates). Their noise correlation thus adds variability orthogonal to the stimulus feature, and therefore has less impact on the performance of a decoder. As a result, reduction of noise correlations between similarly-tuned neurons is an efficient way to improve the precision of a population response, and the decoded information.

To test whether noise correlations between similarly-tuned orientation channels was more suppressed than those for orthogonally-tuned channels, we explored the shape of noise correlations as a function of pairwise tuning dissimilarity. At each timepoint, we fit a linear regression to the baselined noise correlations to test whether stimulus-induced noise correlation was influenced by tuning dissimilarity.

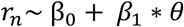

The noise correlation between two channels, *r_n_*, was modelled as a function of their absolute tuning dissimilarity, θ, with an intercept, β_0_, included to capture changes in the mean noise correlation over time. The coefficient for tuning dissimilarity thus describes the extent to which noise correlations vary as a function of tuning dissimilarity, where a positive beta coefficient would describe increase suppression of noise correlations for similarly-tuned channels. This model captured the structure of noise correlations across the 7 possible pairwise tuning dissimilarities as two metrics that we analysed independently.

### Noise correlations and precision

Noise correlations are, by definition, an across-trial metric, reflecting correlated variability in response to repeated presentations of a sensory input, and cannot be measured at the level of a single trial. However, population coding theories suggest that reductions in noise correlations can serve to enhance the information contained in population codes (Averbeck et al., 2006), and empirical evidence supports noise correlation reduction as a mechanism by which attention operates on neuronal activity (Cohen & Maunsell, 2009). Two hypotheses arise: Firstly, greater suppression of mean noise correlations should correlate with higher representational precision. Secondly, that representational precision should be higher when the slope of noise correlation suppression is higher (where noise correlations are more attenuated for similarly-tuned orientation channels.

To test these hypotheses, we independently calculated the across-participant correlation between the precision of the orientation response function and both the mean noise correlation and the slope of noise correlations (β_0_ and β_1_, respectively) estimated from the regression modelling of noise correlations. For each participant, precision was averaged in the time-window used to estimate noise correlations (in steps of 20ms, with a ± 10ms around this), and averaged across both items in the array. This gave an estimate of precision and noise correlations at the same timepoints, per participant. The Pearson’s correlation coefficient was calculated across-participants for each of these time points, giving rise to a time series of the correlation between representational precision and noise correlation parameters (see Figure 4C-D).

To assess significance, we applied a Bonferroni correction based on the number of timepoints in the temporal window of interest (up to 1-second after stimulus onset), comparing the significance of the correlation at each timepoint to this Bonferroni-corrected p-value. This is the same temporal window used for cluster permutation testing in other analyses in this paper. For clarity, we highlight timepoints that reached significance in the unadjusted analysis (Figure 4C-D, pink markers), and were significant after Bonferroni-correction (Figure 4C-D, dark pink markers).

### Statistical Analysis

To interrogate how our derived metrics of tuning curve precision and amplitude related to both the brain and behaviour, we used generalised linear modelling. GLMs were implemented for each participant, modelling the relationship between tuning curve parameters and behavioural and physiological measures across trials. We tested for group significance, at the second-level, by conducting permutation tests across time (to overcome the multiple comparison problem) using the first-level beta coefficients. All permutation tests were conducted using 10,000 permutations, with a cluster forming threshold determined by the one-sided t-value defined by the sample size for the analysis and a significance threshold of 0.05. Permutation tests used a 1-second time-window after stimulus onset to test for effects during early sensory encoding.

To test the hypothesis that the precision with which sensory information is encoded is related to subsequent working-memory recall, we implemented a GLM at each time point of the tuning curve parameter time-series to independently model whether tuning curve precision and amplitude are predictive of subsequent performance. As only one item was probed on each trial, yet we estimate the tuning curve for both items in the array, we only considered the tuning curve of the item that was subsequently probed. We excluded trials with decision times (the time taken to begin the response phase) more than three standard deviations from the within-participant mean and used absolute response error as our behavioural measure. We subtracted the circular mean of the signed response error distribution prior to taking its absolute value, to account for any overall motor biases in orientation reproduction. We standardised (z-scored) the tuning curve parameter of interest at each time point before inclusion as a regressor, and included an intercept term. We conducted one-sided permutation tests to assess significance under the a-priori belief that higher tuning curve amplitude and precision (more information, and higher quality) would be predictive of lower response error across trials.

To test whether variability in baseline physiological markers of attention (alpha power and pupil size) predicted tuning curve precision and amplitude, we conducted a series of GLMs. First, we collapsed our baseline attentional markers over a predefined baseline period (500 ms immediately prior to stimulus presentation). We then tested whether these baseline measures were independently related to the tuning curve parameters (precision and amplitude). Specifically, we independently regressed baseline alpha power onto tuning curve precision, and baseline alpha power onto tuning curve amplitude. Similar models were implemented to test for effects of baseline pupil size.

To test whether baseline alpha power and pupil size were related to one another, we implemented a linear mixed-effects model (LMM) using the *lmerTest* package in R (Kuznetsova et al., 2017). This takes all single trial data across participants and explores whether baseline alpha power is predictive of baseline pupil size at the single-trial level. As baseline pupil size is heavily affected by feedback on the previous trial, we included a fixed effect for previous trial feedback to account for this. We included random effects for previous trial feedback and participant, to account for between-participant differences in the effect of previous-trial feedback and mean pupil size.

For analysis of the EEG data and baseline alpha power, the full 20 participants were included in analysis. For analysis of the relationship with baseline pupil size, 2 participants were excluded due to excessive noise in the epochs, with a final analysed sample of 18 participants for pupillometry analyses. We performed one-sided permutation tests to assess significance, using a cluster forming threshold defined by the one-sided t-value defined by the sample size of the analysis and a significance threshold of 0.05. Specifically, we had three a-priori predictions: Firstly, that high precision of the orientation response function would predict lower working-memory error across trials. Secondly, that lower baseline alpha power (more alpha suppression) over posterior channels would be associated with increased precision. Thirdly, that increased pupil constriction would be related to increased precision.

## Acknowledgements

This research was supported by a Glyn Humphreys Scholarship to S.R.C., a Medical Research Council grant (MR/W01971X/1, MR/L019639/1) and Biotechnology and Biological Sciences Research sLOLA grant (BB/W003392/1) to J.X.O’R., and a BBSRC AFL Fellowship (BB/R010803/1) and European Research Council grant (FORAGINGCORTEX, 101076247) to N.K. The funders had no role in study design, data collection and analysis, decision to publish, or preparation of the manuscript. Views and opinions expressed are those of the author only and no not necessarily reflect those of the funders. Neither the European Union nor the granting authority can be held responsible for them. We wish to thank Kia Nobre, who supervised and funded the original study from which data are taken, and conversations that shaped the conceptual development of this study. For the purpose of Open Access, the author has applied a CC-BY public copyright license to any Author Accepted Manuscript (AAM) version arising from this submission. The authors declare no competing interests.

## Author Contributions

S.R.C. and J.X.O’R. conceived the study. S.R.C. carried out the analysis and wrote all code. S.R.C and J.X.O’R. interpreted the analysis results. S.R.C and J.X.O’R. prepared the manuscript. S.R.C, N.K., and J.X.O’R edited the manuscript. J.X.O’R supervised the research.

