## Supplementary Materials for "Precision of sensory representations in cortex is indexed by alpha power and pupil dilation, and tracks the dynamics of correlated noise"

### Supplementary Information

#### 1 – Dynamic changes in amplitude of the orientation response function

Our two-stage decomposition method for analysing the orientation response function gives rise to two meaningful parameters – the *precision* (quality) and *amplitude* (overall strength) of the orientation response function at each time point.

For completeness, we present a visualisation of the temporal profile of the amplitude parameter (Supplementary Figure 1A). We observed (as for precision) two distinct peaks in the time-course of representational amplitude: a first early peak around 100ms after stimulus onset, and a second later peak around 300-400ms after stimulus onset. Amplitude shares a similar temporal profile to precision during early sensory encoding.

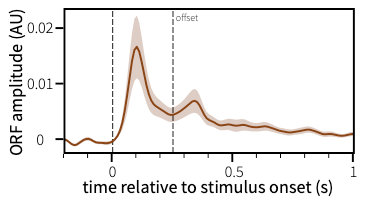

Supplementary Figure 1 -Stimulus onset is associated with rapid increases in the amplitude of sensory information

#### 2 - Amplitude of the orientation response function is unrelated to behavioural performance, baseline alpha power or baseline pupil size

We considered not only whether the precision of the orientation response function (the quality of information) was related to behaviour and baseline biomarkers of attentional state, but also whether the amplitude of this function (the amount of information) was associated with these measures of performance and attention. (A) We found that amplitude of the orientation response function was not significantly associated with later recall error on the working memory task. Amplitude was also not significantly predicted by either pre-stimulus alpha power (B) or pre-stimulus pupil size (C).

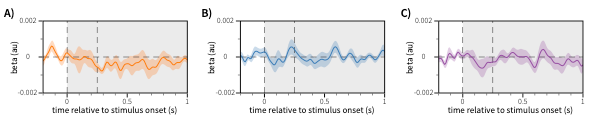

Supplementary Figure 2 - amplitude of the orientation response function is unrelated to working memory error (A), baseline alpha power (B) and baseline pupil size (C)

#### 3 – Formal mediation analysis of effects of baseline alpha power and pupil size on representational precision

To assess whether the effect of alpha power on precision is mediated by pupil size, we conducted a formal mediation analysis using the Baron and Kenny regression framework. In the main article, we observed (using linear mixed-effects modelling) that pre-stimulus pupil size was predictive of pre-stimulus alpha power, and that pupil and alpha both predict neural precision. To determine whether their relationships with neural precision reflected shared or dissociable variance, we fit a regression model that contained both biomarkers simultaneous as independent variables predicting neural precision. We then compared the beta coefficients of the two biomarkers between the solo model (containing only one biomarker) and the combined model (containing both) to determine whether inclusion of the other altered the relationship with neural precision.

We observed no significant difference in the time-course of beta coefficients for either pre-stimulus marker when both were considered in the same model (‘combined model’) vs. their respective independent models (‘solo model’). These results suggest that pre-stimulus alpha power and pupil size – while correlated at the single-trial level – may contribute partially independent variance to trial-by-trial fluctuations in neural precision, arguing against a simple mediation account in which one pre-stimulus marker fully explains the effect of the other.

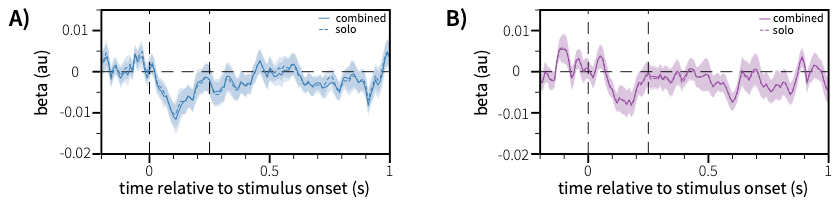

Supplementary Figure 3 – Visualisation of mediation analysis. A) The time-course of the relationship between prestimulus alpha power and sensory precision is shown for the combined model (blue, solid line) and the solo model (blue, dashed line). No significant differences emerged when comparing these two beta coefficient timeseries, suggesting that the inclusion of pupil size as a covariate did not influence the relationship between prestimulus alpha power and sensory precision. B) The time-course of the relationship between prestimulus pupil size and sensory precision is shown for the combined model (purple, solid line) and the solo model (purple, dashed line). No significant differences emerged when comparing these two timeseries, suggesting that inclusion of prestimulus alpha power as a covariate did not alter the relationship between prestimulus pupil size and sensory precision.

#### 4 – Visualisation of raw noise correlations

Here, we show that raw noise correlations are highest for orientation channels with highest tuning similarity (Supplementary Figure 4B), and suppressed in response to stimulus onset (Supplementary Figure 4C). Raw noise correlations are most strongly suppressed for orientation channels with high tuning similarity (Supplementary Figure 4B and 4C, dark blue lines).

In the main article we present visualisation and analysis of baselined noise correlations, such that we explore the change in noise correlations induced by stimulus onset. Baselining was performed to highlight these stimulus-induced changes. As noise correlation suppression also varied as a function of tuning dissimilarity (main article figure 3E), this leads to a ‘knee’ in the raw noise correlations when visualised as a function of tuning dissimilarity (Supplementary Figure 4B, right panel). To consider this interaction when modelling noise correlations as a function of tuning dissimilarity, one would need to fit a model with more free parameters with extra dependencies, resulting in relatively high model complexity when only 6 data points are used to fit the curve. Baselining noise correlations leaves the temporal shape of noise correlation suppression preserved while simply recentring the data. This simplifies the model substantially, allowing a simple linear regression to accurately capture both changes in mean noise correlation (main article figure 3D) and the effect of tuning curve dissimilarity (main article figure 3E).

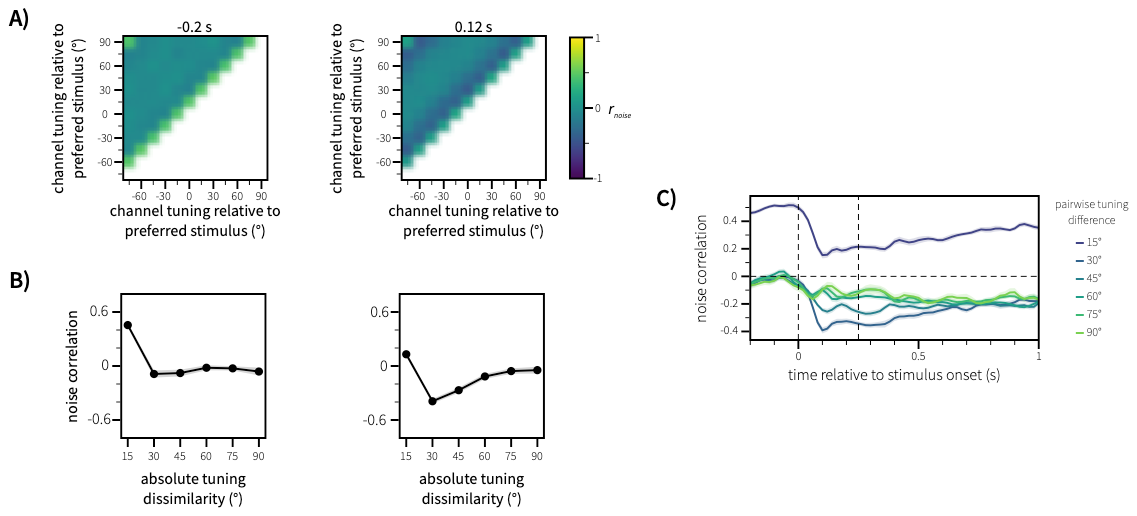

Supplementary Figure 4 – Raw noise correlations between orientation channels. A) Pairwise noise correlations are calculated giving rise to a matrix of noise correlations between orientation channels across time. This is shown for two distinct time periods: during the pre-stimulus period, and during active sensory processing (-0.2s and 0.12s relative to stimulus onset, respectively). B) Raw noise correlations are visualised as a function of the unsigned pairwise tuning difference between orientation channels. C) Raw noise correlations rapidly decrease after stimulus onset for all levels of tuning dissimilarity.

#### 5 - Mahalanobis distance based decoding of orientation information across visual cortical sensors

Previous work on the representational content of scalp EEG activity has used the mahalanobis distance metric to decode orientation from scalp activity (e.g. Wolff et al., 2017; van Ede et al., 2018). Fundamentally this is a pattern-based classifier, where the mahalanobis distance metric measures the multivariate dissimilarity between a test trial and other data. We also implemented a mahalanobis-distance-based pipeline to explore the robustness of our findings across multiple mathematically different, but conceptually similar approaches to orientation decoding. Decoding was implemented independently for each item in the stimulus array, per trial and time point. We describe the implementation here.

To measure the information content of recorded scalp activity, we employed a multivariate decoding approach using the Mahalanobis distance metric to measure the pattern similarity between activity evoked by stimuli of different orientations, using the 17 most posterior sensors. First, the preprocessed, epoched data were lightly smoothed using a gaussian kernel (20ms standard deviation), and the pattern across sensors was demeaned at each time point. Separately for each item in the encoding array, we used a leave-one-out approach to decoding to estimate the orientation preference of the posterior scalp activity at each time point.

First, all training trials were binned based on their orientation relative to the left-out test trial. Bins were created in steps of 4°, and contained trials within ±11 degrees of the centre bin angle, resulting in 45 overlapping bins (Supplementary Figure 5A). We chose this bin width to approximate the half wave rectified cosine kernels used in the population encoding-decoding approach described in the main body of the paper. Comparable results are found using broader orientation bins, and a sparser sampling of the orientation space. The contribution of each individual trial, *t*, to the mean of orientation bin activity was weighted (*w_t_*) by its distance from the bin centre (*θ_b_*) such that while bins were broad and overlapping, each trial had a unique contribution to the activity in each orientation bin.

$$w_{t}=\cos(\frac{\theta_{t}- \theta_{b}}{2* binwidth*\pi})$$

$$\bar{b}_{i}= \frac{\sum(w_{t} \times x_{t})}{\sum w}$$

We then calculated the mahalanobis distance between the weighted mean of activity in each bin and the test trial. We used the pseudo-inverse covariance matrix of all training trials (i.e. excluding the left-out test trial) when calculating the mahalanobis distance. Where a matrix is invertible, the pseudoinverse is identical to the inverse – however, where the matrix is not invertible, the pseudoinverse minimises the least squares problem to estimate the inverse covariance matrix. This results in a mahalanobis distance for each orientation bin, reflecting the pattern similarity in the scalp activity between test trial and the orientation bins – small values reflecting a short distance (high similarity) and large values reflecting a greater dissimilarity in activity. To aid interpretation, we took the negative of these distance values such that the metric now reflects *similarity*, which more positive values reflecting greater similarity between the left-out test trial and the training trials in the feature bin. This allowed us to create, for each trial and time point, a similarity profile, or tuning curve, between the left-out test trial and trials with varying relative orientations. Where there is orientation tuning in the neural signal, this metric will be more positive for similar orientations, and less positive for more dissimilar orientations.

##### Tuning Curve Decomposition

The decomposition approach for the mahalanobis distance-derived tuning curve differs slightly to the approach used for the population encoding-decoding model due to subtle differences in their data.

Estimation of the *precision* of the tuning curve was identical – using a non-linear least squares optimisation routine on the min-max rescaled tuning curve data to fit a flexible cosine model of the form:

$$y_{\theta} \sim\beta_{1}\cos(\alpha\theta)$$

where $\theta$ is the angular distance (in radians) between the centre orientation of the bin and the trial’s stimulus orientation, and *α* indexes the precision of the tuning curve (where larger values indicate narrower, more orientation-selective tuning). We also included a free-parameter for amplitude scaling $\beta_{1}$ to accurately fit the precision parameter.

This fitted tuning curve precision parameter was then lightly smoothed with a gaussian kernel (20ms standard deviation), and this smoothed precision parameter was used to model tuning curve amplitude. In the second stage, we took the similarity values (negative distances) and added the minimum distance value such that all distances were strictly positive. We did this as the cosine function is strictly positive in this feature space, so the tuning curve profile maps more closely onto the model range. The precision parameter at each time point was constrained to have a minimum value of 0.001 here – any lower, and the design matrix vector is a vector of ones, and the model fails to converge.

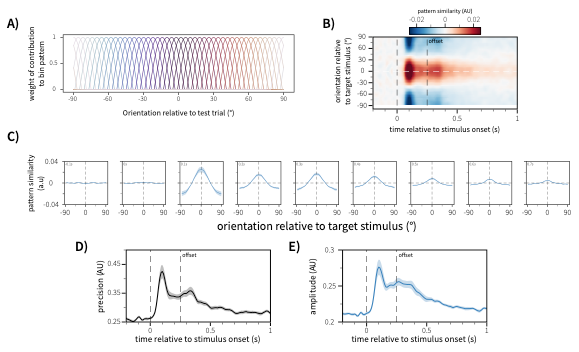

Supplementary Figure 5 – A) structure of the overlapping orientation bins used for mahalanobis distance orientation decoding. Trials were binned based on their relative orientation to the left-out test trial. Individual contributions of each trial to the mean activity of the orientation were cosine-weighted by their angular distance from the centre of the bin. B) Pattern similarities between the left out trial and all other possible orientations are demeaned then visualised to show the stimulus-centred tuning curve from this analytical pipeline. C) The tuning curve is averaged across time windows 50ms either side of the time point shown in the top-left of each sub-panel. This shows the relative amounts of orientation-specific information contained in posterior scalp EEG activity as a function of time. Stimulus onset is associated with a rapid increase in both the precision (D) and amplitude (E) of the orientation information available across these sensors.

For each trial and time point in the epoch of interest, we then fit a generalised linear model (GLM) to fit a minmax-scaled cosine function (of the bin centre orientations) with a shape defined by the previously estimated precision parameter. The beta coefficient of this scaled cosine regressor is thus our best estimate of the amplitude of the tuning curve for a given level of precision.

$$d_{pos} \sim\beta* minmax(\cos\left( \theta_{b}\cdot\alpha\right))$$

Where *d_pos_* is the strictly positive pattern similarity between the test trial and the orientation bins, θ_b_ is the centre orientation of the orientation bin, and *α* reflects the previously estimated value for tuning curve precision for the given trial and timepoint.

##### Results

We first asked when, in the immediate post-stimulus period, the orientation of the stimuli could be decoded from brain activity. As pattern similarity was calculated for bins of varying orientations relative to the test trial, at the high temporal resolution of EEG, we could reconstruct the time resolved orientation function using this mahalanobis distance pipeline (Supplementary Figure 5B). This highlighted an orientation response function that is qualitatively similar to the orientation response function generated through the population encoding-decoding model. We also present the pattern similarity curve across orientations (Supplementary Figure 5C) in the same time-windows as presented for the encoding-decoding approach (main article, Figure 2D), demonstrating the similarities in the timing of decoded orientation information.

We also implemented the decomposition approach, taking the pattern similarity function over relative orientations and modelling it into our two meaningful computational parameters: *precision* (Supplementary Figure 5D) and *amplitude* (Supplementary Figure 5E). Similar to the encoding-decoding model approach, this decomposition method highlights two peaks of precision relative to the onset and offset of the stimulus array.

We also qualitatively replicate the findings of the encoding-decoding model, showing the functional relevance of the precision of this orientation response function. We find that precision as estimated from in this mahalanobis distance orientation decoding pipeline is significantly predictive of working-memory error across trials (Supplementary Figure 6A), where higher precision is associated with lower working-memory error later in the trial. We also find qualitatively similar relationships between precision and baseline alpha power (Supplementary Figure 6B) and baseline pupil size (Supplementary Figure 6C), whereby lower baseline alpha power and a more constricted pupil size are predictive of higher precision during sensory processing.

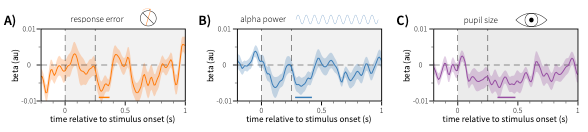

Supplementary Figure 6 - precision of the orientation response function estimated using mahalanobis distance decoding of orientation information is related to working-memory error (A), baseline alpha power (B) and baseline pupil size (C)
